# TADA – a Machine Learning Tool for Functional Annotation based Prioritisation of Putative Pathogenic CNVs

**DOI:** 10.1101/2020.06.30.180711

**Authors:** J. Hertzberg, S. Mundlos, M. Vingron, G. Gallone

**Affiliations:** Max Planck Institute for Molecular Genetics, Ihnestraße 63, 14195 Berlin; Charité Universitätsmedizin Berlin, Charitéplatz 1, 10117 Berlin

**Keywords:** copy-number-variants, structural variants, pathogenicity prediction, functional annotation, TADs, machine-learning

## Abstract

The computational prediction of disease-associated genetic variation is of fundamental importance for the genomics, genetics and clinical research communities. Whereas the mechanisms and disease impact underlying coding single nucleotide polymorphisms (SNPs) and small Insertions/Deletions (InDels) have been the focus of intense study, little is known about the corresponding impact of structural variants (SVs), which are challenging to detect, phase and interpret. Few methods have been developed to prioritise larger chromosomal alterations such as Copy Number Variants (CNVs) based on their pathogenicity. We address this issue with TADA, a method to prioritise pathogenic CNVs through manual filtering and automated classification, based on an extensive catalogue of functional annotation supported by rigorous enrichment analysis. We demonstrate that our machine-learning classifiers for deletions and duplications are able to accurately predict pathogenic CNVs (AUC: 0.8042 and 0.7869, respectively) and produce a well-calibrated pathogenicity score. The combination of enrichment analysis and classifications suggests that prioritisation of pathogenic CNVs based on functional annotation is a promising approach to support clinical diagnostic and to further the understanding of mechanisms that control the disease impact of larger genomic alterations.

## Introduction

The investigation of the genetic causes of rare developmental disorders and, ultimately, the molecular diagnosis of rare disease patients relies on the accurate detection and prioritisation of disease-causing DNA variants. It follows that the accurate identification and prioritisation of candidate disease-associated genetic variation is a fundamental question in human genetic research. The disease impact of single nucleotide polymorphisms (SNPs) and small Insertions and Deletions (InDels) has been the focus of extensive study (Shastry 2002; Montgomery et al. 2013; Wright et al. 2018). Comparatively little is known about the mechanisms and disease impact of structural variants (SVs), including unbalanced SVs, collectively known as Copy Number Variants (CNVs). CNVs have significant and pervasive impact on phenotypic variability and disease: they can affect gene dosage (Huang et al. 2010) and modulate basic mechanisms of gene regulation (Spielmann et al. 2018). In addition, CNVs have been shown to disrupt topologically associating domains (TADs) (Dixon et al. 2012) and can rewire long-range regulatory architectures, resulting in pathogenic phenotypes (Lupiáñez et al. 2015; Kraft et al. 2019).

One of the reasons why CNVs and SVs, in general, are poorly understood is because they are difficult to reliably detect, filter and interpret given current sequencing technology. New experimental approaches such as long-read sequencing (Schadt et al. 2010) combined with novel, long-read specific algorithms for read alignment (Li 2018; Sedlazeck et al. 2018) and SV detection (Cretu Stancu et al. 2017; Sedlazeck et al. 2018; Heller and Vingron 2019) are allowing a more thorough survey of the spectrum of large variation in healthy and diseased human genomes (Collins et al. 2019; Audano et al. 2019). This raises the need for methods to interpret, score and prioritise SVs to support clinical practice.

Ongoing efforts to annotate the potential contribution of SVs to disease suggest the possibility of using functional annotation to stratify SV calls by relevance and/or predicted pathogenic potential (Han et al. 2019). In terms of tools and methods to prioritise pathogenic CNVs, a number of approaches have been proposed (Ganel et al. 2017; Spector and Wiita 2019; Poszewiecka et al. 2018). ClinTAD (Spector and Wiita 2019) and TADeus (Poszewiecka et al. 2018) focus on providing a visual framework to aid a human expert in manually surveying and flagging likely relevant SVs. SVScore (Ganel et al. 2017) aggregates SNP-level CADD (Kircher et al. 2014) scores, integrating single nucleotide-based deleteriousness prediction over the length of a SV. This is based on the assumption that SV effects can be thought of as agglomerates of single-nucleotide-level effects, which is generally unlikely to be the case, given growing evidence of SV impact on, for instance, gene dosage (Huang et al. 2010) and regulatory context (Spielmann et al. 2018). Recently, Kumar et al. 2019 introduced SVFX, a machine learning framework to quantify pathogenicity for somatic and germline CNVs. SVFX represents, to our knowledge, the first flexible machine learning based model for SV pathogenicity prediction. However, the classifier relies on somatic variants as a proxy for pathogenicity as opposed to a set of germline variants annotated as pathogenic. While a subset of these variants is actually pathogenic, the model still likely trains on the differences between somatic and common germline mutations, rather than pathogenic versus non-pathogenic.

Here, we present the TAD annotation tool (TADA), a method to annotate CNVs in the context of their functional environment, based on a rich set of coding as well as non-coding genomic annotation data. The annotation data is centred around TAD boundaries, which serve both as proxy for the regulatory environment (in that they limit the genomic annotation potentially affected by the CNV to the loci between boundaries) and as annotation themselves. TADA is designed to assess the functional relevance of user-specified input sets of CNVs of unknown clinical relevance by one of two methods: a) annotation, followed by manual filtering, or b) machine-learning based automated classification. Importantly, our machine learning models for duplications and deletions are trained on a set of annotated pathogenic variants (DECIPHER, (Firth et al. 2009)) and rigorously driven by functional evidence: we carefully assess the biological relevance of each of the annotations considered by performing enrichment tests, comparing the expected and observed overlap of pathogenic versus non-pathogenic CNVs and functional annotation data. We demonstrate feasibility of our approach on two separate test sets: 1) a set of ClinVar variants and 2) and a split of DECIPHER and non-pathogenic variants not included in our training data, resulting in an AUC for the deletion model of 0.8125 and 0.8897, respectively. Both the deletion and the duplication model are able to place more than 50% of pathogenic variants among a set of rare non-pathogenic variants in the first ten ranks of 100 based on predicted pathogenicity. TADA is available free-of-charge under the MIT license and can be customised for prioritising or classifying CNVs from different disease contexts.

## Results

### Enrichment Analysis of Pathogenic CNVs

Here, we performed a comprehensive enrichment analysis of the pathogenic DECIPHER variant data set (Firth et al. 2009) in comparison to a curated set of common, and therefore unlikely pathogenic, CNV calls (MacDonald et al. 2014; Aguirre et al. 2019; Collins et al. 2019; Audano et al. 2019). Our purpose was to assess whether we could identify contrasting patterns of enrichment/depletion in a pathogenic set with respect to a control set. We reasoned that, if this was the case, the discriminating annotations would be excellent candidates for a feature set of a classifier to distinguish pathogenic from non-pathogenic variants. In our analysis, we account for size differences between the variant sets (Supplementary S1) and for the non-uniform mutation rate across the genome (Audano et al. 2019) (Methods for details) (which would have artificially inflated fold changes) by building GC-content isochores (Costantini et al. 2006) and constraining bootstrapping with bins of comparable GC-content signal. The set of annotations tested in the enrichment analyses was based on evidence from Collins et al. 2019 and Audano et al. 2019 including coding and non-coding annotation as well as conservation and predicted loss-of-function (pLoF) metrics. We additionally integrated TAD boundaries (Dixon et al. 2015), CTCF bindings sites (Dunham et al. 2012), genes associated with developmental disease (DDG2P) as well as genes predicted to be haplosufficient (HS Genes) and haploinsufficient (HI Genes) (Huang et al. 2010). The results for pathogenic and non-pathogenic deletions are shown in Fig. 1.

**Figure 1.**
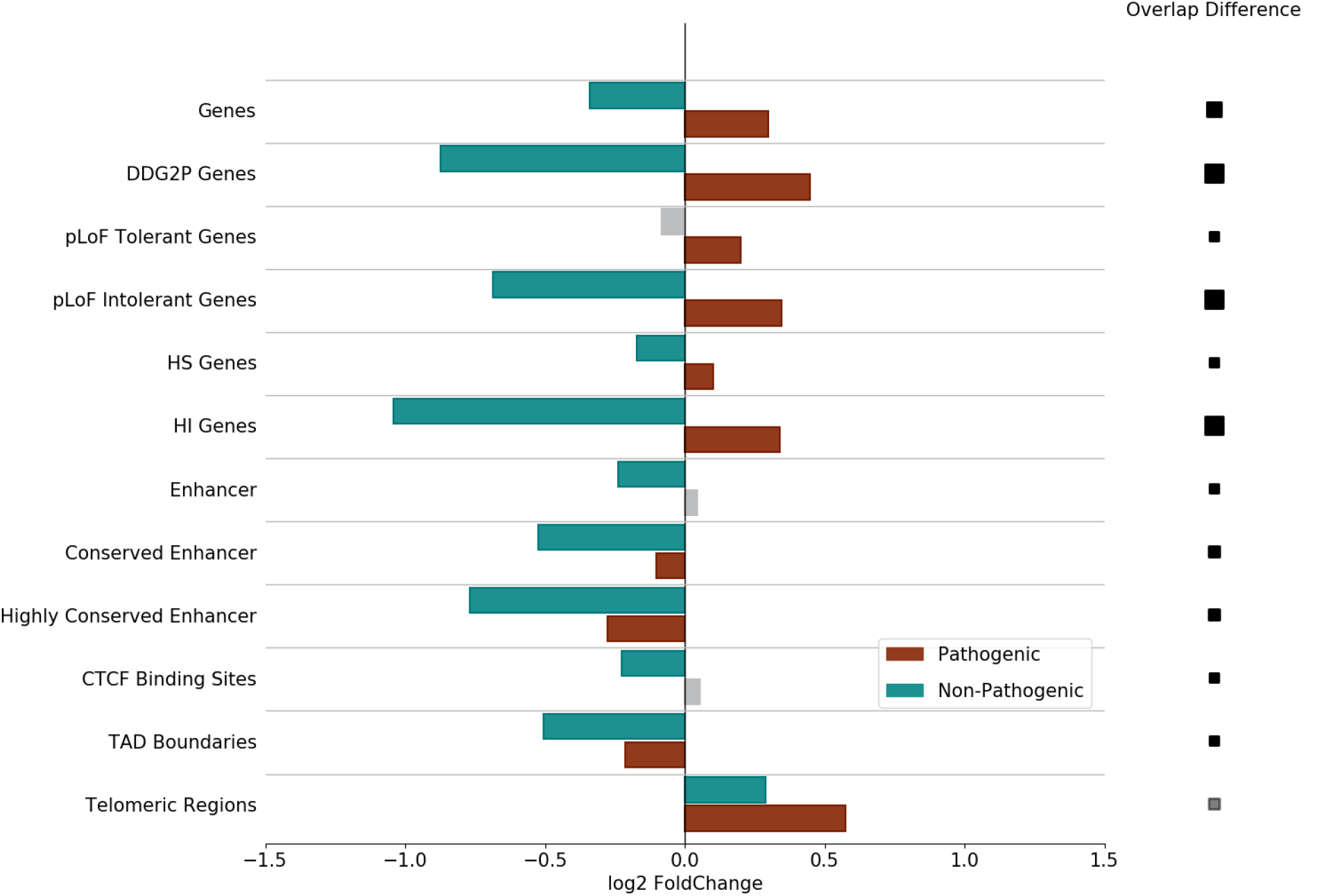
Enrichment Analysis of non-pathogenic and pathogenic deletions. The figure shows the log_2_(fold change) for expected and observed variant overlap for each set of genomic annotations based on 10, 000 simulations. The size of the squares on the right side of the figure is proportional to the overlap FC difference between pathogenic and non-pathogenic deletions. Grey bars and squares indicate a non-significant FC (*q*-value ≤ 0.01).

We conducted our enrichment tests within the genome association tester (GAT) framework (Heger et al. 2013), a bootstrap-based method to test for enrichment or depletion of genomic segments in background annotations accounting for a variety of confounding factors (Methods for details). Briefly, we generated a number of randomly distributed, size-matched genomic segments in each simulation and computed overlaps with sets of genomic annotation. We computed overlaps over all simulations (*expected* overlaps, see Methods) and compared them to *observed* overlaps, producing fold-change (FC) values and associated significance for each annotation set. To account for potentially diverging patterns of enrichment/depletion due to the variant type, we ran enrichment tests separately for deletions and duplications using 10, 000 simulations.

In agreement with what was previously shown by Collins et al. we observe significant depletion of non-pathogenic deletions in coding regions (logFC = −0.340, *q*-val = .15 * 10^−3^) and regulatory regions (logFC = −0.240, *q*-val = .15 * 10^−3^). The depletion signal increases with predicted haploinsufficiency (logFC = −1.041, *q*-val = .15 * 10^−3^) of the affected coding regions and conservation of the regulatory regions (logFC = −0.768, *q*-val = .15 * 10^−3^). While we observe a strong significant depletion of non-pathogenic deletions in pLoF intolerant genes, we do not detect significant depletion in pLoF tolerant genes. We observe stronger enrichment in pLoF intolerant genes with respect to background gene annotation (logFC = −0.686, ‥67 * 10^−3^), confirming previous observations reporting increased depletion of common structural deletions in coding regions intolerant to LoF-mutations. In agreement with Audano et al. 2019 we observe significant enrichment of non-pathogenic deletions in extended (See Methods for an definition of *extended*) telomeric regions (logFC = 0.289, *q*-val = .8727 * 10^−3^). Additionally, our combined set of non-pathogenic deletions is significantly depleted in TAD boundaries (logFC = −0.506, *q*-val = .15*10^−3^) and CTCF binding sites (logFC = −0.109, *q*-val = .72*10^−3^).

The enrichment analysis of pathogenic DECIPHER deletions reveals patterns of significant enrichment in all functional annotation except FANTOM5 enhancer regions, TAD boundaries and extended telomeric regions. The pathogenic deletions are significantly enriched in coding regions (logFC = 0.316, *q*-val = .15 * 10^−3^), with increased enrichment for DDG2P genes (logFC = 0.447, *q*-val = .15 * 10^−3^). We observe increased enrichment in pLoF intolerant genes (logFC = 0.346, *q*-val = .8 * 10^−3^) compared to pLof tolerant genes (logFC = 0.208, *q*-val = .15 * 10^−3^) as well as HI genes (logFC = 0.319, *q*-val = .15 * 10^−3^) compared to HS genes (logFC = 0.122, *q*-val = .13 * 10^−3^). The pathogenic deletions are also significantly enriched in extended telomeric regions (logFC = 0.594, *q*-value = .15 * 10^−3^). We are not able to detect significant enrichment of our pathogenic set in any set of the enhancer annotations, regardless of the extent of sequence conservation. Instead, we observe significant depletion of pathogenic deletions in highly conserved enhancers (logFC = −0.272, *q*-val = .15 * 10^−3^). Interestingly, the enrichment analysis also reveals a significant depletion of pathogenic deletions in TAD boundaries (logFC = −0.188, *q*-val = .15 * 10^−3^). The analysis of duplications reveals similar patterns of enrichment for pathogenic and non-pathogenic variants (Supplementary Fig. S3).

While the set of functional annotation we introduced above shows potential as a feature set to distinguish between pathogenic and non-pathogenic variants, it is not likely to represent the full spectrum of regulatory activity across the genome. Doan et al. 2016 observed an enrichment of CNVs impacting human accelerated regions (HARs) (Pollard et al. 2006) i.e. regions that are highly conserved across vertebrates with increased divergence in humans, in individuals with autism spectrum disorder (ASD). This suggests potential brain associated regulatory function of HARs (Pollard et al. 2006). To test a wider range of genomic annotation such as HARs as distinguishing features, we set out to conduct further enrichment analyses. Motivated by evidence of CNV enrichment in segmental duplications (SDs) (Kim et al. 2008), we included SDs in our analysis. Given the increasing evidence for the impact of non-coding variation in Mendelian disorders, localising in highly conserved, tissue-specific active distal regulatory elements such as enhancers (Short et al. 2018), we also included ChromHMM annotations (Ernst and Kellis 2012). The enrichment results of SDs, HARs and ChromHMM annotation for deletions and duplications are shown in Supplementary Fig. S4 and S5, respectively. Both non-pathogenic and pathogenic deletions are significantly depleted in polycomb-repressed regions (logFC = −0.477, *q*-val = .7 * 10^−3^, logFC = −0.168, *q*-val = .14 * 10^−2^, respectively) and HARs (logFC = −0.399, *q*-val = .1 * 10^−3^, logFC = −0.216, *q*-val = .1 * 10^−3^, respectively). We observe no significant depletion or enrichment of pathogenic or non-pathogenic deletions in any other ChromHMM annotation or in SDs. In contrast we observe significant enrichment of non-pathogenic duplications in SDs (logFC = 0.375, *q*-val = .1 * 10^−3^). We also observe significant depletion of non-pathogenic duplications in polycomb-repressed regions (logFC = −0.248, *q*-val = .1 * 10^−3^) and small but significant depletion in HARs (logFC = −0.560, *q*-val = .1 * 10^−3^).

TADs are known to approximately represent windows of constrained regulatory interactions (Shen et al. 2012). We reasoned that for TADs of high regulatory relevance, pathogenic CNVs are likely depleted across the entire TAD environment due to their potential effect on the corresponding regulatory context. We therefore set out to investigate the enrichment of pathogenic CNVs in TADs stratified by their regulatory importance. We assumed that the regulatory importance of a TAD can be approximated by the conservation of enhancer annotation and the pLoF intolerance of coding annotation within the TAD environment (Methods for details). We henceforth refer to the resulting set of TAD annotations as *TAD-centric* annotations. Supplementary Fig. S6 and S7 show the results of the TAD-centric enrichment analysis for, respectively, deletions and duplications. We observe significant enrichment of non-pathogenic deletions and duplications in TADs lacking known coding or regulatory annotation (logFC = 1.128, *q*-val = .47*10^−3^, logFC = 1.436, *q*-val = .15*10^−3^, respectivley) and significant depletion of non-pathogenic CNVs in most TADs containing coding or regulatory elements. In contrast, we observe significant enrichment of pathogenic deletions in TADs containing coding and regulatory annotation with an increased enrichment in TADs encompassing at least one highly loF intolerant gene (logFC = 0.240, *q*-val = .28 * 10^−3^). However, we do not detect a signal of enrichment or depletion for pathogenic duplications in TAD-centric annotations and cannot confirm increased enrichment of pathogenic deletions in TADs containing highly conserved enhancers compared to TADs with less conserved enhancers. Taken together, the results point towards selective pressure towards deletions based on the entire affected regulatory domain rather than individual coding and non-coding annotation.

### TADA

We used the observed patterns of enrichment and depletion to inform feature selection in our TAD-annotation tool. However, we are well aware that the relevance of the selected features for the prioritisation of putative pathogenic variants will differ, based on disease and sample context. To account for the variable relevance of features we allowed user-defined annotation alongside the default feature set driven by the results of the enrichment analysis. A schematic and a detailed description of the TADA workflow is shown in Fig. 2.

**Figure 2.**
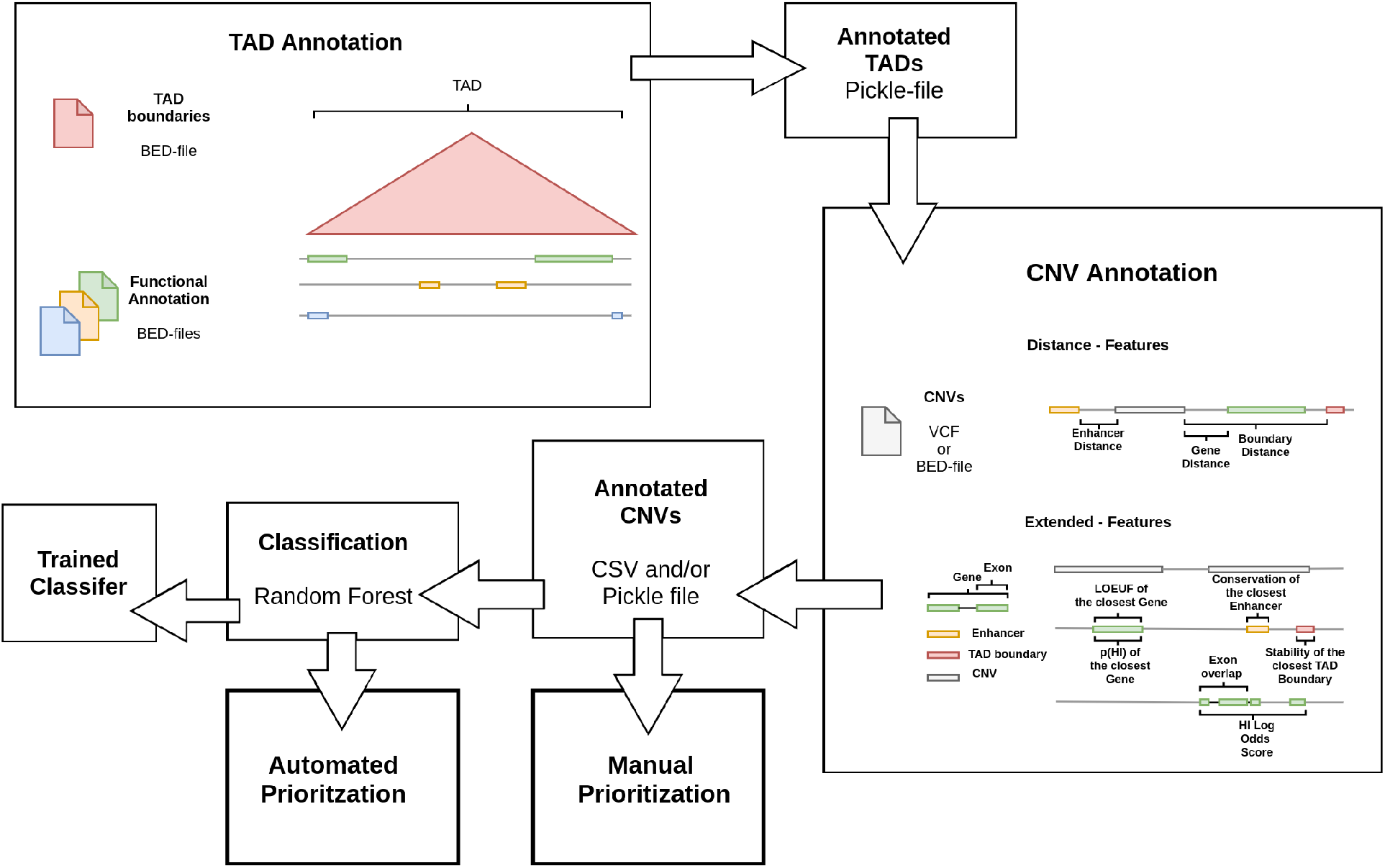
Generalised Workflow of the TADA tool. The basis for the CNV annotation are BED-files of TAD-boundaries and any additional genomic annotation. In a first step, the annotation sets are sorted into the corresponding TAD environment based on genomic position. The resulting annotated TAD regions are used as a proxy of the regulatory environment during the CNV annotation. The feature set for the annotation process consists of *Distance* features, which describe the distance to the closest element of an annotation set in the same TAD environment, and *Extended* features, which refer to metrics describing the functional relevance of the corresponding coding or regulatory element. The annotated CNVs can either be manually prioritised or classified by a model trained to distinguish between pathogenic and non-pathogenic variants. Depending on the user input the annotation is either based on a predefined feature set that includes *Extended* and *Distance* features or on individual *Distance* feature set derived from the user defined set of genomic annotations.

### Pathogenicity Prediction

We demonstrate the viability of functional annotation as basis for the prioritisation of putative pathogenic CNVs by training classifiers using the TADA tool and evaluating their predictive performance. We split the variant set used for the enrichment analyses into deletions and duplications and trained separate random forest classifiers on a total of 14 functional annotation derived features (Table S2). The features included: distance to the closest gene, FANTOM5 enhancer, CTCF binding site and TAD boundary in the corresponding TAD environment. Additionally, we included the loss-of-function observed/expected upper bound fraction (LOEUF) (Collins et al. 2019) and Haploinsufficiency potential (Huang et al. 2010) for the closest gene as well as evolutionary conservation of the closest enhancer. Finally, we used the HI log Odds score (Huang et al. 2010), the TAD stability of the closest TAD boundary (McArthur and Capra 2020) and a feature corresponding to the overlap of a CNV with potential regulatory regions based on pcHi-C (Jung et al. 2019). For each variant type, we split our original data into training and test set (70/30) stratified by label distribution and trained a random forest classifier. We then evaluated the performance of the parameter-tuned classifiers with 5-fold cross-validation, the test set split of our original data and on a set of pathogenic and benign ClinVar deletions and duplications without overlap to our training data. This resulted in AUC values for the deletion model of 0.8340 (5-CV), 0.8897 (ClinVar) and 0.8125 (Test-Set). The AUC values for the duplication model are 0.8100 (5-CV), 0.8473 (ClinVar) and 0.7907 (Test-Set split) (Fig 3 **C**).

**Figure 3.**
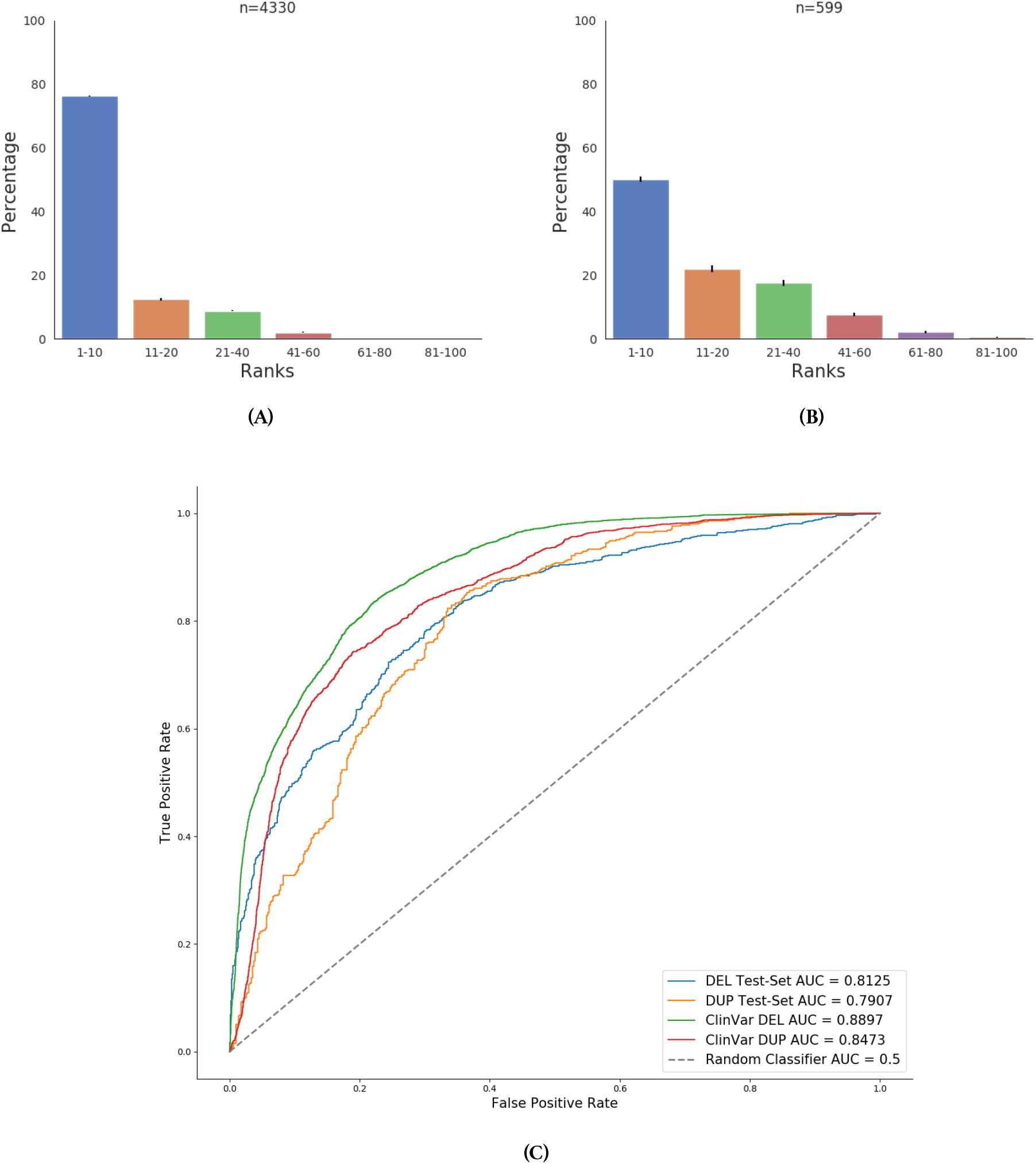
Predictive Performance of the Deletion and Duplication Classifiers. **A** and **B** shows the ranking performance of the deletion and duplication classifiers, respectively. We binned the ranks into groups of ten and computed the percentage of variants placed among these ranks. **C** shows the ROC-Curves and AUC values for the deletion and duplication classifiers based on the separate test-set and ClinVar variants.

Our analysis of the predictive performance across multiple test sets provides an indication of the classifier’s ability to distinguish between pathogenic and non-pathogenic variant given a hard threshold on the pathogenicity probability i.e. the probability of the variant to belong to the pathogenic class. In clinical practice a common scenario is to distinguish a single pathogenic variant from a set of rare non-pathogenic variants. We assumed that a criteria for classifier usability should be as follows: the pathogenic variant should be placed among the first ten ranks based on predicted pathogenicity. For this purpose the classifiers predicted class probabilities need to be *calibrated* i.e. the proportion of true positives needs to be close to the pathogenic probability for a given probability threshold (Zadrozny and Elkan 2001). Since our classifiers seemed to be well calibrated based on fraction of positives compared to the mean predicted value (Supplementary Fig. S8), we set out to analyse its *ranking* ability. We generated size-matched batches of 100 deletions or duplications, containing a single pathogenic ClinVar and 99 rare variants GnomAD variants (Methods for details). For each batch we computed the rank of the pathogenic variants based on predicted pathogenicity and obtained the standard deviation of individual ranks. The ranking performance of our classifiers for both deletions and duplications is shown in Fig. 3 **A**/**B**. The deletion classifier places from 4, 330 batches 76% of the pathogenic deletions among the first ten ranks and a cumulative 97% among the first 40. From a total of 599 batches the duplication classifier places 50% of pathogenic variants among the first ten ranks and cumulatively 90% among the first 40.

Our classifier performs well on developmentally associated pathogenic variants and shows a high test-set and ranking performance for the ClinVar database, indicating that the model can be applied to a disease and sample context unrelated to the training set. However, both DECIPHER and ClinVar focus mainly on the coding effect of variants which is likely to be reflected in our trained classifier. We therefore decided to determine the most relevant features and identify potential biases in our trained model. To account for any biases introduced by correlated features, we clustered the features based on partial correlation and computed the mean loss in accuracy after permutation of highly correlated feature clusters (Strobl et al. 2007) (Methods for details). The mean-loss and standard deviation across 30 computations with different random seeds is shown in Fig. 4. As expected, the results indicate that our trained model relies mainly on coding rather than non-coding functional annotation. The most relevant features for the trained classifier are the predicted haploinsufficiency of the closest gene and the HI Log Odds score, followed by the distance to the closest gene and its intolerance to LoF-mutations. Regulatory annotations are of lower importance for classification. This is in agreement with our enrichment results, where we could not observe enrichment of pathogenic DECIPHER variants in FANTOM5 enhancer annotations. We expect regulatory annotation such as enhancers to increase in importance as the number of pathogenic training non-coding variants increase.

**Figure 4.**
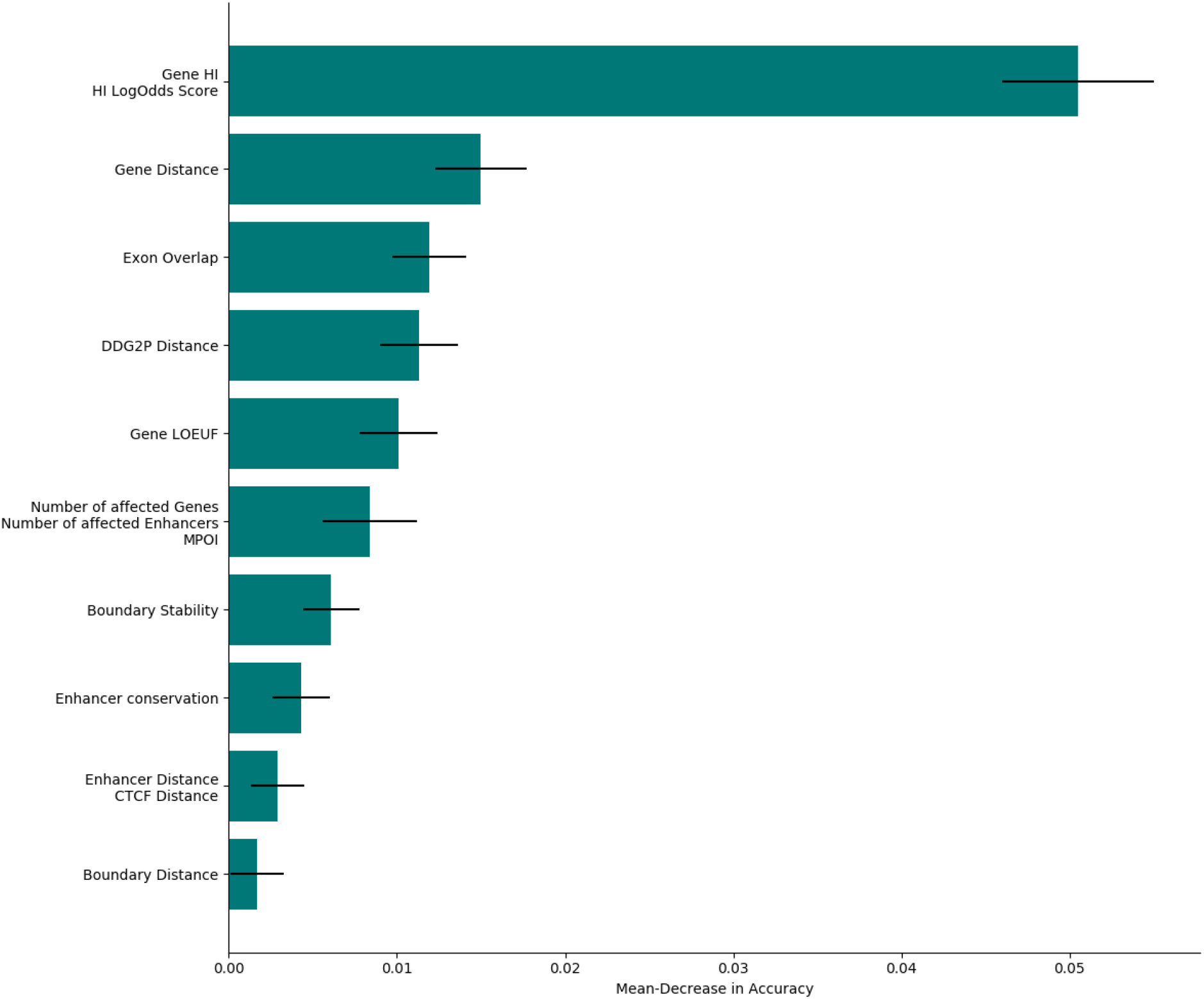
Feature Importance of the Deletion Model. The figure shows the mean loss in accuracy after permutation of highly correlated feature clusters (see Methods for a detailed description of individual features). The standard deviation based on 30 sampling runs with variable random seed is indicated by black lines.

## Discussion

The prioritisation and identification of disease-causing genomic variants is an active field of genomic research. SNPs and InDels and their relation to human disease have been the focus of intense study (Shastry 2002; Montgomery et al. 2013). Although early evidence suggests SVs are a major contributor to genome-wide variation, comparatively little is known about their disease impact at an individual and population level. This can be attributed to technical limitations, namely the accurate identification of SVs and precise SV breakpoint calling. It follows that few studies have, so far, focused on the prioritisation and ranking of large sets of SV calls to highlight smaller and more manageable subsets of SVs with higher predicted disease potential.

Recent experimental as well as algorithmic advancements in the field of SV detection are leading to an increase in publicly available catalogues of SVs (Collins et al. 2019; Audano et al. 2019). These catalogues are focused mainly on common i.e. likely non-pathogenic variants and reveal patterns of depletion in coding and regulatory functional annotation. Due to the lack of publicly available data, an analysis on a comparatively comprehensive catalogue of pathogenic SVs is missing.

In this work, we used the DECIPHER database of pathogenic CNVs to analyse the potential of functional annotation and SV overlap as a distinguishing feature between pathogenic and non-pathogenic CNVs. We conducted an extensive series of enrichment tests, identifying contrasting patterns of enrichment between pathogenic and non-pathogenic CNVs for multiple sets of genomic annotation. We observed significant depletion of non-pathogenic variants in coding and regulatory regions, positively correlated with the intolerance to LoF-mutations and predicted haploinsufficiency of coding regions as well as the primary sequence conservation of regulatory regions. In our analyses we observed a contrasting pattern of enrichment for pathogenic variants in coding regions and CTCF binding sites, providing additional evidence for the potential of functional annotation as a feature to identify pathogenic variants. We conducted a second enrichment analysis with a wider range of non-coding associated annotations. The analysis included ChromHMM annotations, SDs and HARs. We were not able to observe a contrasting pattern of enrichment and depletion in this group of annotations, suggesting that ChromHMM annotations, SDs and HARs do not represent genomic regions with differential selective pressure towards perturbating variation. In a final enrichment analysis we investigated the overlap of pathogenic and non-pathogenic CNVs with TAD-centric features i.e. TADs stratified by regulatory importance. We observed a significant depletion of non-pathogenic variants in TADs containing regulatory and coding annotation. In contrast, pathogenic variants are significantly enriched in TADs of regulatory importance. This suggests that the selection towards variation affecting functional annotation is likely to extend to entire regulatory domains.

The enrichment analyses also revealed significant depletion of pathogenic variants in TAD boundaries and highly conserved enhancers, indicating that regulatory functional annotation is less important than coding sequence-centric annotation in aiding the discrimination of pathogenic versus non-pathogenic variants based on the data sets used in these analyses. This is perhaps unexpected, given there is increasing evidence on the role of variants impacting non coding regulatory elements on rare disease (Spielmann et al. 2018; Short et al. 2018). Accounting for the focus of DECIPHER on the coding rather than non-coding effect of CNVs, we reasoned that the significant depletion of pathogenic variants in non-coding functional associated annotation is a consequence of investigator bias in our set of annotated pathogenic variants. We anticipate that, with larger less biased pathogenic SV repositories becoming available, observed genome-wide SV impact on regulation will yield a stronger signal. Current prioritisation methods should therefore provide the flexibility to account for both coding and non-coding effects.

For this purpose we developed TADA, a flexible annotation method to prioritise pathogenic CNVs based on their overlap with functional annotation. The tool offers the option to manually *prioritise* variants i.e. it returns the annotated CNVs as a list that can be sorted by each of the annotations. It also allows for machine-learning-based prioritisation using a random-forest model trained on a functional annotation based feature set. The default feature set that we provide includes coding and non-coding associated features, motivated by the results of our enrichment analysis. Alternatively, the user can provide a set of genomic annotation associated with a disease or sample context to generate an custom feature set. Even though, based on the enrichment analysis, non-coding features such as TAD boundaries and enhancer conservation do not assist in the differentiation of pathogenic from non-pathogenic variants, we decided to include them in the default feature set. By doing this, we accounted for the investigator bias in the DECIPHER catalogue of pathogenic variants. We further used the default feature set to train random forest classifiers for deletions and duplications and evaluated their predictive performance by measuring the AUC and found they performed well on a separate test set and ClinVar variants. We assessed the precision of the pathogenicity score over batches of rare variants combined with a single pathogenic variant. Both the deletion and duplication model performed well, placing more than 60% of the pathogenic variants among the first ten rank based on the pathogenicity score. This indicates that the pathogenicity score is a close approximation of true pathogenic effect in our test set. As expected, the analyses of the classifier’s feature importance revealed dependency on coding regions rather than regulatory regions, mirroring the results of our enrichment analyses. We therefore recommend the application of the automated prioritisation using the pre-trained random forest model with focus on the coding rather than non-coding effect of CNVs. Since TADA is trained on the DECIPHER variants, which were identified as the cause of developmentally associated disease phenotypes, we cannot guarantee that the pre-trained model is able to accurately classify variants in a different disease context. Hence, we provide the possibility to manually prioritise variants based on a user-defined or on the default feature set, which also includes features accounting for the non-coding effect of CNVs.

Due to recent experimental and algorithmic approaches SVs can be reliably identified and their role in clinical diagnostics is beginning to be established. Still, the proportion of balanced SVs in publicly available databases of pathogenic variation is comparatively low, limiting any approach focusing on the prioritisation of pathogenic variants, including TADA. to unbalanced SVs. Nevertheless, our results show that the prioritisation of pathogenic CNVs based on functional annotation is a promising approach. With the likely increasing number of more comprehensive available variant catalogues, we aim to improve the predictive power of our classifier and adapt our approach to other classes of disease-relevant genomic structural variation.

## Materials and Methods

### Variant Sets

We obtained 20, 990 CNVs from DECIPHER (Firth et al. 2009). We filtered the CNVs according to their pathogenicity and size, and chose to retain variants categorised as *pathogenic*, *likely pathogenic* or *unknown* with a size larger than 50*bp*. Since DECIPHER serves as database to analyse candidate i.e. potential disease-causing variants, we reasoned that a large proportion of variants with *unknown* are still likely pathogenic. We noticed that multiple DECIPHER calls were overlapping and possibly representing the same variant, we selected the smallest variant for each pair/cluster of overlapping variants based on a 90% reciprocal overlap. Finally, we only kept variants located on autosomes.

The common i.e. *non-pathogenic* variant set is a compendium built from four different data sources: a recent publication by the Eichler group featuring SVs identified via deep PacBio sequencing of 15 individuals (Audano et al. 2019), a collection of approximately 14, 891 SVs published by the GnomAD consortium (Collins et al. 2019), a set of CNVs called from the UK Biobank data set (Aguirre et al. 2019) and CNVs obtained from the Database of Genomic Variants (DGV) (MacDonald et al. 2014). All variants were either already mapped to GRCh37 or converted to GRCh37 using the UCSC *LiftOver* tool (Kent et al. 2002). In order to match the pathogenic variants we limited the set of non-pathogenic variants to CNVs. We discarded rare and potentially deleterious variants by applying individual filters to each of the data sources: we filtered for *Shared* or *Major* variants published by the Eichler group for i.e. variants present in all or ≥ 50% of the samples, GnomAD SV variants with an allele frequency (AF) *>* 0.1, UK Biobank variants supported by 3 or more samples and DGV variant reported in more than one publication. Of these variants we only kept non-pathogenic located on autosomes larger than 50*bp*. To account for overlapping variants between different sources of non-pathogenic variants, we clustered variants with a reciprocal overlap greater or equal to 90%. For each pair/cluster of overlapping variants we selected a single variant based on their origin, with the following prioritisation order: Audano et al., GnomAD-SV, DGV and UK Biobank. We reasoned that variant calls reported based on sequencing – especially long-read-sequencing – provide more precisely resolved breakpoints than variants reported based on array-CGH or SNP-arrays.

While the pathogenic variants were called using array-CGH, the GnomAD and Audano variants are based on WGS and long-read-sequencing, respectively. The difference in experimental methods is reflected in the size distribution of pathogenic and non-pathogenic variants (Supplementary Fig. S1). To account for the size bias across variants sets, we binned the non-pathogenic variants by size using an empirical cumulative distribution function (ECDF) with bin size 60. We then sampled for each bin the same number of pathogenic variants. For bins with a higher number of non-pathogenic than pathogenic variants, we sampled the same number of non-pathogenic variants as pathogenic variants without replacement. This resulted in a size and number matched variant set of 6, 128 deletions and 3, 476 duplications. The proportion of non-pathogenic variants by data source changed during the size-matching, due to the lack of short pathogenic variants. Supplementary Fig. S2 shows the proportion of variants by pathogenicity and data source before and after the size matching.

For the analysis of the classifiers’ ranking performance, we used GnomAD CNVs with an AF <= 0.1 as rare benign variants. We filtered for duplicated variants in a similar fashion as the training set i.e. searching for clusters of overlapping variants. Since the pathogenic ClinVar were significantly larger, we picked the largest variants of each cluster, to maximise the number of variants after size-matching. Then we discarded any CNVs overlapping with our training data (90% reciprocal overlap).

We obtained ClinVar CNVs from () on October 24, 2019 restricting the search by Type of variation to copy number gain, copy number loss, deletions and duplications. First, we stratified the variants by type of variant i.e. deletions and duplications. Then, we separated both deletions and duplications into pathogenic (Pathogenic and Likely pathogenic annotation) and benign (Benign and Likely benign annotation) and only kept variants located on autosomes. Finally, we discarded any duplicated variants (as described above for the DECIPHER dataset) and those overlapping with our training data (90% reciprocal overlap) in each set of pathogenic/non-pathogenic deletions and duplications.

### Annotations

We obtained hg19 TAD boundaries from human embryonic cells (Dixon et al. 2012) as described by McArthur and Capra 2020 annotated with *boundary stability*.

For both the enrichment testing and the training of the pathogenicity classifier we used FANTOM5 enhancer annotations (Andersson et al. 2014) without any tissue specifications. We annotated the enhancers using aggregated PhastCons and PhyloP scores (Siepel et al. 2005) i.e. the mean of base-wise conservation scores over the enhancer interval. The *conserved* and *highly conserved* enhancers in the enrichment analysis correspond to FANTOM5 enhancers with aggregated PhastCons scores over the 75% and 90% percentiles given the background distribution over all enhancer annotations.

We obtained gene annotations from the GnomAD consortium and exon annotations for the computation of the exon overlap feature from GENCODE (Harrow et al. 2012). We stratified the gene annotations by predicted Haploinsufficiency (*p*(*HI*)) (Huang et al. 2010) and intolerance to loss-of-function mutations i.e. LOEUF (Karczewski et al. 2019). For the enrichment analysis we defined genes with *p*(*HI*) *>* 0.9 and *p*(*HI*) < 0.1 as *HI Genes* and *HS Genes*, repsecitvly. Genes with LOEUF< 0.1 and LOEUF*>* 0.9 as *ploF Tolerant Genes* and *ploF Intolerant Genes*.

We obtained hg19 CTCF binding site annotations from ENCODE i.e. irreproducible discovery rate (IDR) optimal ChIP-seq peaks (ENCODE Accession Number: ENCSR000EFI).

The computation of overlap with potential regulatory regions identified in pcHi-C data is based on data from Jung et al. 2019. We selected the *P-O-interactions* (*p*-value=3) for each gene contained in our set of annotated genes and computed the overlap of a CNVs with each interacting fragment i.e. 1 if the CNV overlaps with a fragment, 0 otherwise. Finally, we divided the sum over all interacting fragment for each gene by the genes LOEUF value.

We obtained hg19 telomeric regions from the UCSC genome browser (Kent et al. 2002) and extended them by 5*mbp* to match the annotations described by Audano et al. 2019.

### Enrichment Testing

We performed the enrichment tests using the *gat-run* protocol of the *Genome Association Tester* (GAT) (Heger et al. 2013). GAT is a bootstrap-based approach used to test the association between sets of genomic intervals. The gat-run.py protocol merges the segments of interest i.e. the CNVs in a preprocessing step, which resulted at first in a coverage (track density) above 90% for each variant set. To avoid false positive enrichment due to size bias, we therefore used the size-matched CNVs sets and reduced the track density for pathogenic and non-pathogenic variants to 30.1% and 22.7%, respectively.

CNVs are non-randomly distributed across the genome. Regions with an increased amount of paralogous repeats such as segmental duplications are prone to non-allelic homologous recombination, a recombination process that can lead to the formation of deletions or duplications (Bailey et al. 2002). To account for the non-random distribution of CNVs, we used GC-content families (Costantini et al. 2006) to split the genome into *isochores*. For each isochore we performed a separate enrichment test using the isochores function of GAT. We used gat-run.py with the following settings: -nbuckets 10000 -bucket-size=960 -num-samples=10000 -counter=segment-overlap. To compare the FCs between pathogenic and non-pathogenic variants we used gat-compare.py.

### Classification and Preprocessing

We annotated the pathogenic and non-pathogenic CNVs with a total of 14 features (Supplementary Table S1). In preparation to the classification we split the data into training and test set (70/30 split). We then imputed missing data in both sets with the mean of the corresponding feature in the training set. To account for the differences in data ranges of raw feature values and decrease the convergence time during training, we scaled all features to a range between 0 and 1. Similar to the imputation process, we fit the scaler on the training data and applied it to the test data. Finally, we trained a Random Forest on the imputed and scaled training set. We then evaluated the performance based on stratified 5-fold Cross-Validation and on the separate test set using AUC.

### Ranking Performance

In order to test the ranking performance of our trained model i.e. its ability to differentiate the true pathogenic variant from a set of rare putative non-pathogenic variants we used the above described rare variants obtained from Collins et al. 2019 and the pathogenic ClinVar variants. We predicted the pathogenicity score i.e. the probability to belong to the pathogenic class using our pretrained random forest classifiers for both rare and pathogenic variants. We then binned the rare variants by size using an ECDF with 60 bins. For each pathogenic variant we selected the closest bin of non-pathogenic variant by size discarding any variants larger than the largest non-pathogenic variants. If the corresponding bin contained 99 or more variants we randomly sampled 99 non-pathogenic variants. We then sorted the non-pathogenic variants and the single pathogenic variant by predicted pathogenicity and used the index in the sorted list as predicted rank. In order to generate standard deviations for the variant placement we repeated the sampling of non-pathogenic variants over 30 (numpy.random.choice) random seeds.

### Feature Importance

We employed hierarchical clustering using the scipy python package to generate clusters of highly correlated features based on the training set of annotated size-matched pathogenic and non-pathogenic CNVs. For each cluster with a maximal distance of one we permutated the correlated feature columns in our training data and computed the predicted accuracy using the pre-trained random forest model. We then reported the difference between the accuracy based on the original and permutated data set. Both accuracies are based on the out-of-bag samples of the random forest model. Using the numpy.random.choice function we generated 30 random seeds between 0 and 100 and repeated the permutation process for each random seed. We then computed the mean and the standard deviation for the distribution of accuracy differences of each cluster.

## Data Access

We provide a table containing download links and obtained dates for both the variant and annotation sets in the supplement (Table S2). The TADA tool including a full documentation, pretrained models for deletions and duplications, genomic annotation underlying our feature set and the respective scripts for preprocessing can be accessed at https://github.com/jakob-he/TADA.

## Acknowledgements

We thank Verena Heinrich, Robert Schöpflin, Uirá Souto Melo, David Heller and Emel Comak for useful discussions. We also thank James Priest and Matthew Aguirre for providing us with access to the CNVs calls from the UK Biobank. This study makes use of data generated by the DECIPHER Consortium. A full list of centers who contributed to the generation of the data is available from http://decipher.sanger.ac.uk and via email from. Funding for the DECIPHER project was provided by the Welcome Trust.

## Disclosure Declaration

The authors declare no competing interests.

## References

Aguirre M, Rivas M, and Priest J. 2019. Phenome-wide burden of copy Number variation in UK Biobank. BioRxiv. 545996.

Andersson R, Gebhard C, Miguel-Escalada I, Hoof I, Bornholdt J, Boyd M, Chen Y, Zhao X, Schmidl C, Suzuki T, et al. 2014. An atlas of active enhancers across human cell types and tissues. Nature. 507: 455.

Audano PA, Sulovari A, Graves-Lindsay TA, Cantsilieris S, Sorensen M, Welch AE, Dougherty ML, Nelson BJ, Shah A, Dutcher SK, et al. 2019. Characterizing the major structural variant alleles of the human genome. Cell. 176: 663–675.

Bailey JA, Gu Z, Clark RA, Reinert K, Samonte RV, Schwartz S, Adams MD, Myers EW, Li PW, and Eichler EE. 2002. Recent segmental duplications in the human genome. Science. 297: 1003–1007.

Collins RL, Brand H, Karczewski KJ, Zhao X, Alföldi J, Khera AV, Francioli LC, Gauthier LD, Wang H, Watts NA, et al. 2019. An open resource of structural variation for medical and population genetics. BioRxiv. 578674.

Costantini M, Clay O, Auletta F, and Bernardi G. 2006. An isochore map of human chromosomes. Genome research. 16: 536–541.

Cretu Stancu M et al. 2017. Mapping and phasing of structural variation in patient genomes using nanopore sequencing. Nature Communications. 8: 1326.

Dixon JR, Jung I, Selvaraj S, Shen Y, Antosiewicz-Bourget JE, Lee AY, Ye Z, Kim A, Rajagopal N, Xie W, et al. 2015. Chromatin architecture reorganization during stem cell differentiation. Nature. 518: 331.

Dixon JR, Selvaraj S, Yue F, Kim A, Li Y, Shen Y, Hu M, Liu JS, and Ren B. 2012. Topological domains in mammalian genomes identified by analysis of chromatin interactions. Nature. 485: 376.

Doan RN, Bae BI, Cubelos B, Chang C, Hossain AA, Al-Saad S, Mukaddes NM, Oner O, Al-Saffar M, Balkhy S, et al. 2016. Mutations in human accelerated regions disrupt cognition and social behavior. Cell. 167: 341–354.

Dunham I, Birney E, Lajoie BR, Sanyal A, Dong X, Greven M, Lin X, Wang J, Whitfield TW, Zhuang J, et al. 2012. An integrated encyclopedia of DNA elements in the human genome.

Ernst J and Kellis M. 2012. ChromHMM: automating chromatin-state discovery and characterization. Nature methods. 9: 215.

Firth HV, Richards SM, Bevan AP, Clayton S, Corpas M, Rajan D, Van Vooren S, Moreau Y, Pettett RM, and Carter NP. 2009. DECIPHER: database of chromosomal imbalance and phenotype in humans using Ensembl resources. The American Journal of Human Genetics. 84: 524–533.

Ganel L, Abel HJ, Consortium F, and Hall IM. 2017. SVScore: an impact prediction tool for structural variation. Bioinformatics. 33: 1083–1085.

Han L et al. 2019. Functional annotation of rare structural variation in the human brain. bioRxiv.

Harrow J, Frankish A, Gonzalez JM, Tapanari E, Diekhans M, Kokocinski F, Aken BL, Barrell D, Zadissa A, Searle S, et al. 2012. GENCODE: the reference human genome annotation for The ENCODE Project. Genome research. 22: 1760–1774.

Heger A, Webber C, Goodson M, Ponting CP, and Lunter G. 2013. GAT: a simulation framework for testing the association of genomic intervals. Bioinformatics. 29: 2046–2048.

Heller D and Vingron M. 2019. SVIM: structural variant identification using mapped long reads. Bioinformatics. 35: 2907–2915.

Huang N, Lee I, Marcotte EM, and Hurles ME. 2010. Characterising and predicting haploinsufficiency in the human genome. PLoS genetics. 6: e1001154.

Jung I, Schmitt A, Diao Y, Lee AJ, Liu T, Yang D, Tan C, Eom J, Chan M, Chee S, et al. 2019. A compendium of promoter-centered long-range chromatin interactions in the human genome. Nature genetics. 1–8.

Karczewski KJ, Francioli LC, Tiao G, Cummings BB, Alföldi J, Wang Q, Collins RL, Laricchia KM, Ganna A, Birnbaum DP, et al. 2019. Variation across 141,456 human exomes and genomes reveals the spectrum of loss-of-function intolerance across human protein-coding genes. BioRxiv. 531210.

Kent WJ, Sugnet CW, Furey TS, Roskin KM, Pringle TH, Zahler AM, and Haussler D. 2002. The human genome browser at UCSC. Genome research. 12: 996–1006.

Kim PM, Lam HY, Urban AE, Korbel JO, Affourtit J, Grubert F, Chen X, Weissman S, Snyder M, and Gerstein MB. 2008. Analysis of copy Number variants and segmental duplications in the human genome: Evidence for a change in the process of formation in recent evolutionary history. Genome research. 18: 1865–1874.

Kircher M, Witten DM, Jain P, O’Roak BJ, Cooper GM, and Shendure J. 2014. A general framework for estimating the relative pathogenicity of human genetic variants. Nature genetics. 46: 310.

Kraft K et al. 2019. Serial genomic inversions induce tissue-specific architectural stripes, gene misexpression and congenital malformations. Nature Cell Biology. 21: 305–310.

Kumar S, Harmanci A, Vytheeswaran J, and Gerstein MB. 2019. SVFX: a machine-learning framework to quantify the pathogenicity of structural variants. bioRxiv.

Li H. 2018. Minimap2: pairwise alignment for nucleotide sequences. Bioinformatics. 34: 3094–3100.

Lupiáñez DG, Kraft K, Heinrich V, Krawitz P, Brancati F, Klopocki E, Horn D, Kayserili H, Opitz JM, Laxova R, et al. 2015. Disruptions of topological chromatin domains cause pathogenic rewiring of gene-enhancer interactions. Cell. 161: 1012–1025.

MacDonald JR, Ziman R, Yuen RK, Feuk L, and Scherer SW. 2014. The Database of Genomic Variants: a curated collection of structural variation in the human genome. Nucleic acids research. 42: D986–D992.

McArthur E and Capra JA. 2020. Topologically associating domain (TAD) boundaries stable across diverse cell types are evolutionarily constrained and enriched for heritability. bioRxiv.

Montgomery SB et al. 2013. The origin, evolution, and functional impact of short insertion–deletion variants identified in 179 human genomes. Genome Research. 23: 749–761.

Pollard KS, Salama SR, King B, Kern AD, Dreszer T, Katzman S, Siepel A, Pedersen JS, Bejerano G, Baertsch R, et al. 2006. Forces shaping the fastest evolving regions in the human genome. PLoS genetics. 2:

Poszewiecka B, Stankiewicz P, Gambin T, and Gambin A 2018. TADeus-a tool for clinical interpretation of structural variants modifying chromatin organization. In: 2018 IEEE International Conference on Bioinformatics and Biomedicine (BIBM), pp. 84–87.

Schadt EE, Turner S, and Kasarskis A. 2010. A window into third-generation sequencing. Human Molecular Genetics. 19: R227–R240.

Sedlazeck FJ, Rescheneder P, Smolka M, Fang H, Nattestad M, Haeseler A von, and Schatz MC. 2018. Accurate detection of complex structural variations using single-molecule sequencing. Nature Methods. 15: 461–468.

Shastry BS. 2002. SNP alleles in human disease and evolution. Journal of human genetics. 47: 561.

Shen Y, Yue F, McCleary DF, Ye Z, Edsall L, Kuan S, Wagner U, Dixon J, Lee L, Lobanenkov VV, et al. 2012. A map of the cis-regulatory sequences in the mouse genome. Nature. 488: 116–120.

Short PJ, McRae JF, Gallone G, Sifrim A, Won H, Geschwind DH, et al. 2018. De novo mutations in regulatory elements in neurodevelopmental disorders. Nature. 555: 611.

Siepel A, Bejerano G, Pedersen JS, Hinrichs AS, Hou M, Rosenbloom K, Clawson H, Spieth J, Hillier LW, Richards S, et al. 2005. Evolutionarily conserved elements in vertebrate, insect, worm, and yeast genomes. Genome research. 15: 1034–1050.

Spector JD and Wiita AP. 2019. ClinTAD: a tool for copy Number variant interpretation in the context of topologically associated domains. Journal of human genetics. 64: 437.

Spielmann M, Lupiáñez DG, and Mundlos S. 2018. Structural variation in the 3D genome. Nature Reviews Genetics. 19: 453–467.

Strobl C, Boulesteix AL, Zeileis A, and Hothorn T. 2007. Bias in random forest variable importance measures: Illustrations, sources and a solution. BMC bioinformatics. 8: 25.

Wright CF et al. 2018. Making new genetic diagnoses with old data: iterative reanalysis and reporting from genome-wide data in 1,133 families with developmental disorders. Genetics in Medicine. 20: 1216–1223.

Zadrozny B and Elkan C 2001. Obtaining calibrated probability estimates from decision trees and naive Bayesian classifiers. In: Icml. Vol. 1. Citeseer, pp. 609–616.

